# Genetic alteration of human *MYH6* is mimicked by SARS-CoV-2 polyprotein: mapping viral variants of cardiac interest

**DOI:** 10.1101/2021.11.23.469709

**Authors:** Praveen Anand, Patrick J. Lenehan, Michiel Niesen, Unice Yoo, Dhruti Patwardhan, Marcelo Montorzi, AJ Venkatakrishnan, Venky Soundararajan

## Abstract

Acute cardiac injury has been observed in a subset of COVID-19 patients, but the molecular basis for this clinical phenotype is unknown. It has been hypothesized that molecular mimicry may play a role in triggering an autoimmune inflammatory reaction in some individuals after SARS-CoV-2 infection. Here we investigate if linear peptides contained in proteins that are primarily expressed in the heart also occur in the SARS-CoV-2 proteome. Specifically, we compared the library of 136,704 8-mer peptides from 144 human proteins (including splicing variants) to 9,926 8-mers from all 17 viral proteins in the reference SARS-CoV-2 proteome. No 8-mers were exactly identical between the reference human proteome and the reference SARS-CoV-2 proteome. However, there were 45 8-mers that differed by only one amino acid when compared to the reference SARS-CoV-2 proteome. Interestingly, analysis of protein-coding mutations from 141,456 individuals showed that one of these 8-mers from the SARS-CoV-2 Replicase polyprotein 1a/1ab (KIALKGGK) is identical to a *MYH6* peptide encoded by the c.5410C>A (Q1804K) genetic variation, which has been observed at low prevalence in Africans/African Americans (0.08%), East Asians (0.3%), South Asians (0.06%) and Latino/Admixed Americans (0.003%). Furthermore, analysis of 4.85 million SARS-CoV-2 genomes from over 200 countries shows that viral evolution has already resulted in 20 additional 8-mer peptides that are identical to human heart-enriched proteins encoded by reference sequences or genetic variants. Whether such mimicry contributes to cardiac inflammation during or after COVID-19 illness warrants further experimental evaluation. We suggest that SARS-CoV-2 variants harboring peptides identical to human cardiac proteins should be investigated as ‘viral variants of cardiac interest’.

## Introduction

Cardiac injury is a prevalent complication associated with COVID-19.^1^ In a study of 100 recently recovered COVID-19 patients, cardiovascular magnetic resonance imaging revealed cardiac involvement or ongoing myocardial inflammation in 78 and 60 patients, respectively.^2^ In another study of 39 consecutive autopsies from patients who died of COVID-19, viral RNA was detectable in the heart of 24 (62%) patients. A large nationwide study from Israel reported that SARS-CoV-2 infection is associated with increased rates of myocarditis, arrhythmia, myocardial infarction, and pericarditis.^3^ Myocarditis has also been reported in a small fraction of individuals after receiving an mRNA COVID-19 vaccine.^4–6^

Despite these phenotypic associations, the mechanisms underlying myocardial inflammation in the setting of COVID-19 infection and vaccination remain unclear. A prevalent hypothesis, known as molecular mimicry, posits that T lymphocytes and/or antibodies that recognize SARS-CoV-2 antigens and mediate virus neutralization may also cross-react against host cardiac proteins and trigger an autoimmune response against cardiomyocytes.^7^ This mechanism has also been suggested to contribute to other inflammatory conditions seen in the context of COVID-19 infection.^8–10^ Indeed, autoimmune sequelae of other infectious diseases have been attributed to mimicry between host and microbial antigens.^11–16^

Advances in next-generation sequencing technologies have facilitated the rapid development of large-scale multi-omic datasets and genomic epidemiology resources to better understand the COVID-19 pandemic. Bulk and single cell RNA-sequencing datasets have elucidated the transcriptional signatures of most healthy human tissues and cell types.^17–20^ Amino acid sequences of human proteins, including genetic variants and immunologic epitopes, are available in UniProt^21^, gnomAD^22^, and Immune Epitope Database (IEDB)^23^. The GISAID database currently hosts 4.85 million SARS-CoV-2 genomes from more than 200 countries.^24^ The availability of such genome-scale data enables us to investigate the potential for molecular mimicry between SARS-CoV-2 and human cardiac proteins.

Here we present a systematic comparison of peptides from human cardiac proteins and SARS-CoV-2 proteins. We show that no 8-mer peptides are identical between the reference sequences of these two groups of proteins. However, when including human and viral genetic variants in this comparison, we found 21 8-mer peptides to be identical between human cardiac proteins and SARS-CoV-2 proteins. Among these, a human genetic variant of *MYH6* (c.5410C>A; Q1804K) is identical to a peptide of the reference SARS-CoV-2 replicase polyprotein. Finally, we propose that the SARS-CoV-2 variants that have peptides identical to human cardiac proteins should be studied as potential ‘viral variants of cardiac interest’.

## Results

### Identification of genes that are overexpressed in cardiac tissue

To identify heart-enriched proteins, we compared the expression of all human protein-coding genes in heart samples (n = 861) versus all non-striated muscle samples (n = 15,718) from the Genotype Tissue Expression (GTEx) project. There were 137 genes expressed at least 5-fold higher in the heart with a Cohen’s D value greater than or equal to 0.5 (**Figure 1, Supplementary Table 1**). Similarly, we compared the expression of all genes between cardiomyocytes (n = 8.9K cells) and non-cardiomyocytes (n = ∼2.5 million cells) across a database of 52 single-cell RNA-sequencing studies covering 62 tissues.^18^ There were 46 genes overexpressed in cardiomyocytes based on the same criteria outlined above (**Figure 1, Supplementary Table 1**). Combining the lists of genes identified from bulk and single-cell RNA-sequencing analyses, we identified a total of 144 candidate cardiac proteins.

**Figure 1.**
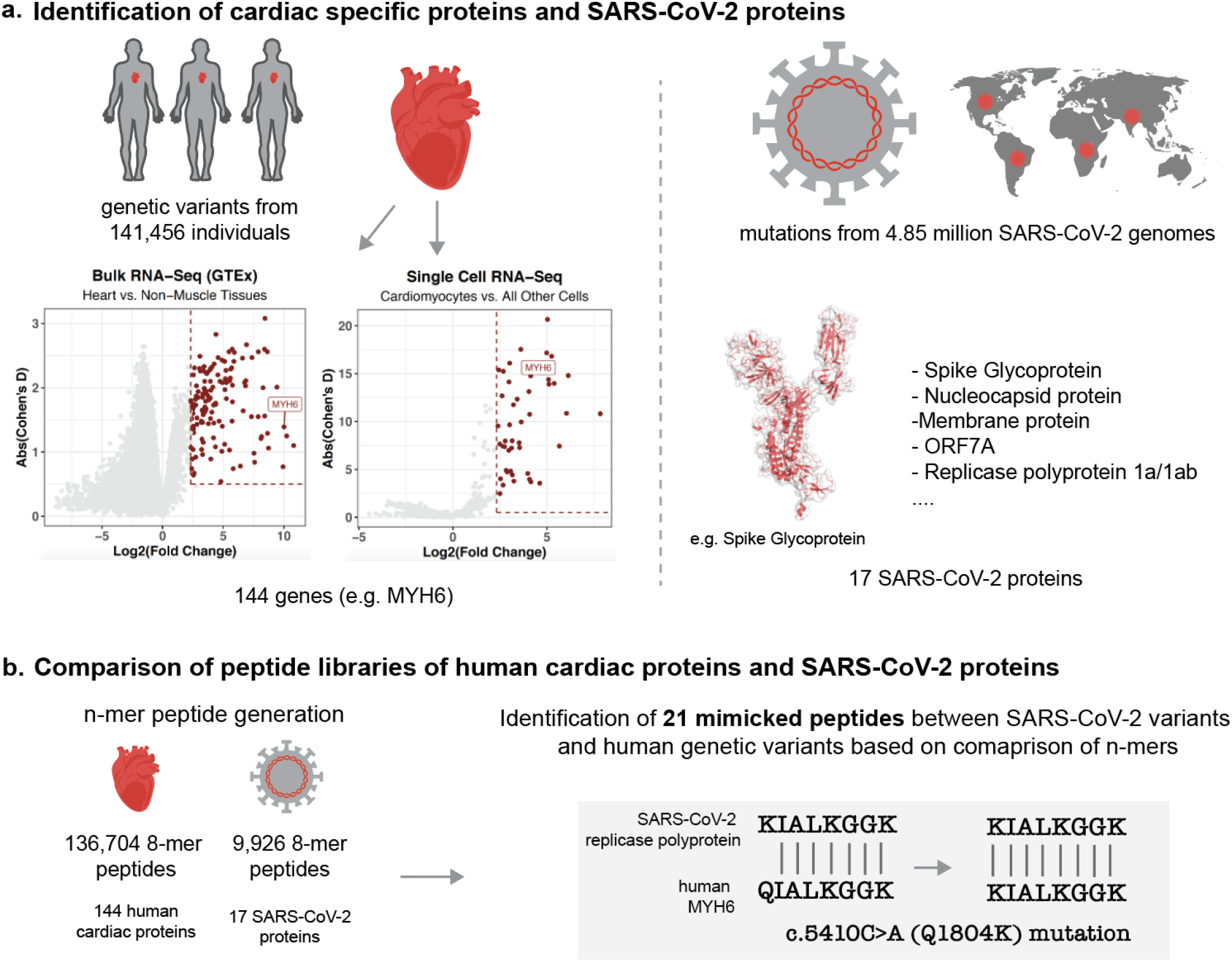
Identification of mimicked peptides between SARS-CoV-2 and human proteins. **a**. Identification of cardiac specific proteins based on analysis of bulk RNAseq and single-cell RNAseq data and identification of SARS-CoV-2 proteins. **b**. Comparison of peptide libraries of human cardiac proteins and SARS-CoV-2 proteins.

### *MYH6* variant is mimicked by an epitope of SARS-CoV-2 Replicase polyprotein 1a/1ab

We computed peptide 8-mers from the reference sequences of the 144 cardiac proteins (including isoforms) and all 17 proteins from the reference SARS-CoV-2 sequence. We then systematically compared the pairwise sequence identity (using Hamming Distance; see **Methods**) for the 136,704 cardiac protein 8-mers with the 9,926 8-mers derived from the SARS-CoV-2 proteins. No peptides were fully identical between these two groups. However, 45 8-mers were nearly identical, with only a single mismatched amino acid (**Table 1**). To determine whether human genetic variation results in any 8-mers which exactly match the reference SARS-CoV-2 proteome, we then analyzed amino acid mutations from 141,456 individuals using the gnomAD database (including 83,623 mutations in the 144 cardiac proteins).^22^

**Table 1:**
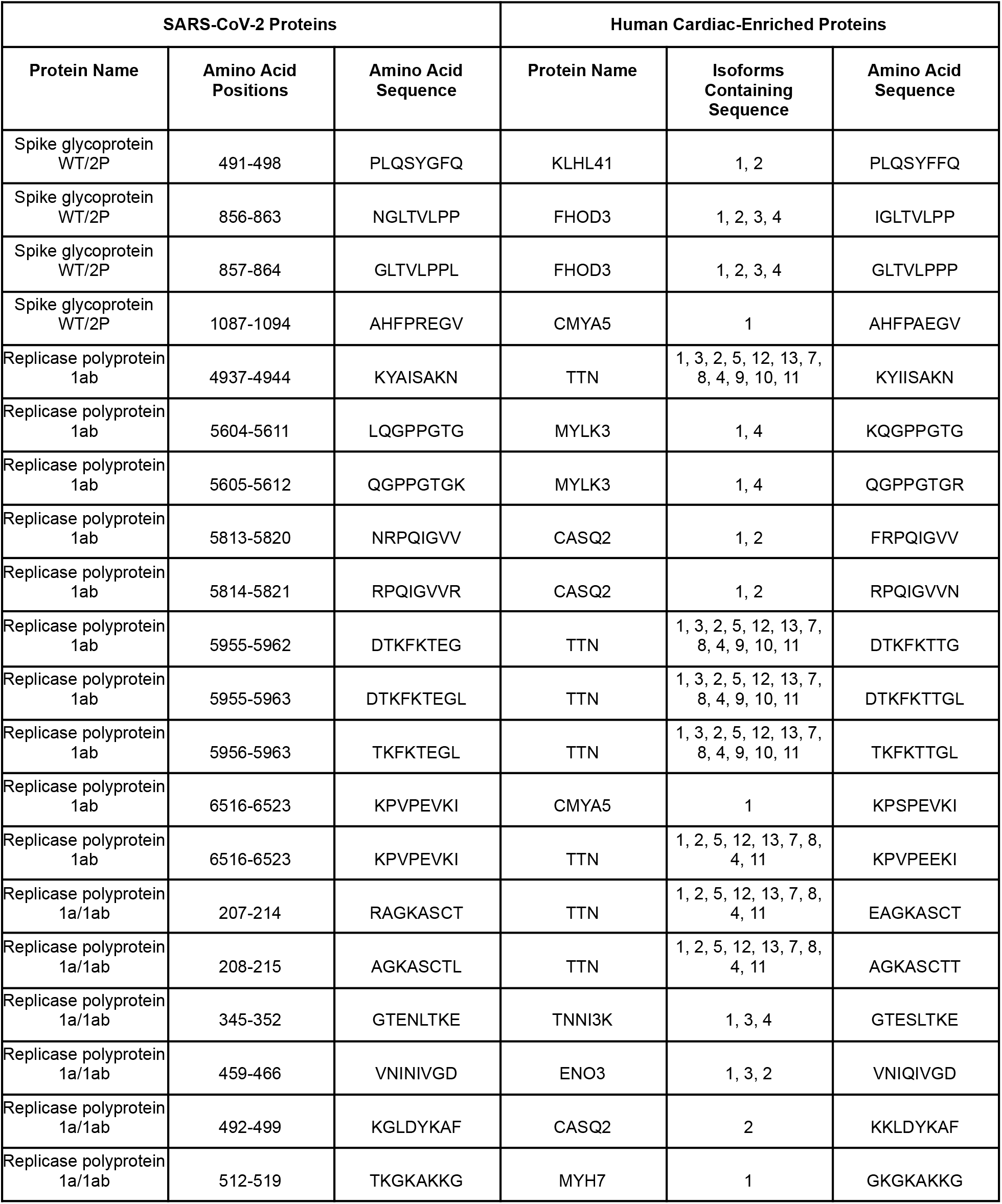
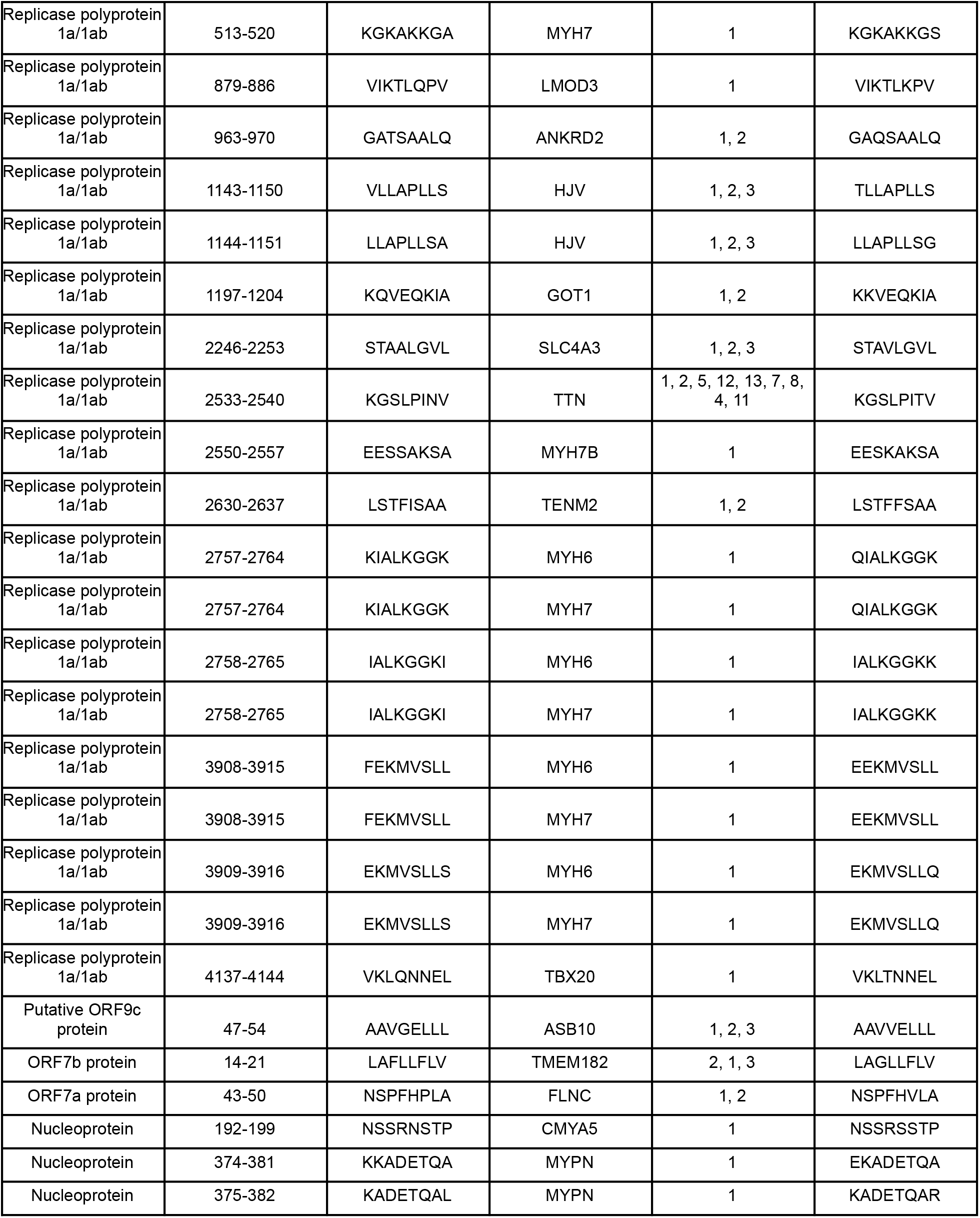
List of peptide pairs from SARS-CoV-2 proteins and human cardiac proteins that have a Hamming distance less than or equal to 1.

Interestingly, one specific 8-mer from the SARS-CoV-2 Replicase polyprotein 1a/1ab (KIALKGGK) is identical to a mutant peptide encoded by the c.5410C>A(Gln1804Lys) variation in human *MYH6* (**K**IALKGGK), which is a subunit of a cardiac motor protein. This genetic variation has been identified in Africans/African Americans (0.08% prevalence), East Asians (0.3% prevalence), South Asians (0.06% prevalence) and Latino/Admixed Americans (0.003% prevalence).^22^ Analysis of peptides from IEDB shows that the non-mutated 7-mer in this peptide (IALKGGK) also overlaps with a known B-cell epitope from SARS-CoV-2 (IALKGGKIVNNWLKQ)^25^. Whether the mimicry between SARS-CoV-2 Replicase polyprotein 1a/1ab and wild-type or mutant *MYH6* contributes to cardiac inflammation in the setting of COVID-19 warrants further investigation.

### SARS-CoV-2 variants harbor epitopes that are identical to peptides of cardiac proteins

We next examined whether SARS-CoV-2 evolution has given rise to variants that harbor peptides identical to human cardiac proteins. To this end, we analyzed 4.85 million SARS-CoV-2 genomes from over 200 countries obtained from the GISAID database.^24^ We identified 21 8-mer peptides from SARS-CoV-2 variants that are identical to cardiac proteins encoded by reference sequences or genetic variants (**Tables 2 and 3**). Of these 21 peptides, the one present in the highest number of viral sequences was STAVLGVL, which mapped to the NSP3 protein (part of Replicase polyprotein sequence) in 4,501 (0.09%) SARS-CoV-2 genomes. This sequence is present in a transmembrane helix of all three described isoforms of the cardiac protein SLC4A3, an anion exchange protein that is associated with short QT syndrome (**Tables 2 and 3**).^26,27^ Although no epitopes containing this full sequence were present in IEDB, a BLAST search at 90% similarity identified a similar epitope (HEAQ**AVLGVL**L) that is reported to bind to both HLA-B*40:1 and HLA-B*58:01.^25,28^ We additionally analyzed the temporal and geographical emergence of the five variant SARS-CoV-2 8-mers that are identical to cardiac protein 8-mers and that occur in over 100 reported SARS-CoV-2 genomes (**Supplementary Figure S1**). Variants with exactly matching 8-mers have occurred sporadically in different geographic locations, but they have not persisted over time (**Supplementary Figure S1**).

**Table 2:**
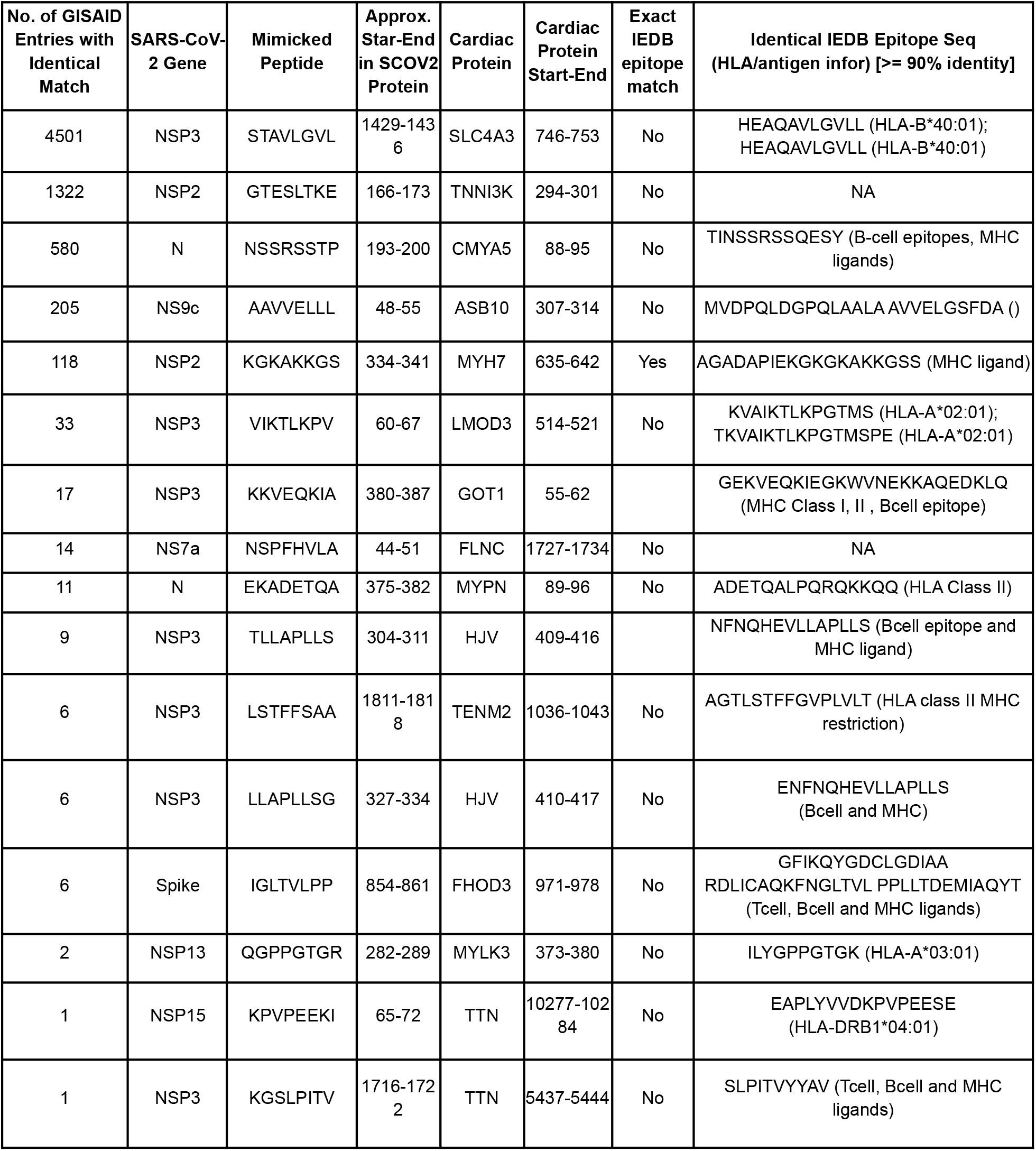
List of identical cardiac peptides found in SARS-CoV-2 variants in GISAID. The NSP proteins are cleaved products of the replicase polyprotein.

**Table 3.** List of mutated cardiac peptide nmers from human genetic variants identical to SARS-CoV-2 variants. The NSP proteins are cleaved products of the replicase polyprotein.

| Human Cardiac Gene | rsID | Mutation consequence | Mutated cardiac Peptide | Wild-type cardiac peptide | No. of GISAID genomes | SARS-CoV-2 gene | IEDB epitope Exact match | IEDB epitope info (>=90 % Seq Identity) |
| --- | --- | --- | --- | --- | --- | --- | --- | --- |
| FLNC | rs374848954 | p.Val1732Leu | NSPFHLLA | NSPFHVLA | 1661 | NS7a | Yes | NSPFH (HLA-A*01:01) |
| LMOD3 | rs370869958 | p.Lys519Arg | VIKTLRPV | VIKTLKPV | 85 | NSP3 | No | NA |
| TMEM182 | rs774398171 | p.Gly215Val | LAVLLFLV | LAGLLFLV | 21 | NS7b | No | NA |
| MYLK3 | rs771870674 | p.Arg380Cys | QGPPGTGC | QGPPGTGR | 1 | NSP13 | No | GPPGTGKSHFAIGLA (B cell, Tcell, MHC ligand) |

## Discussion

Understanding the mechanistic basis of acute cardiac injury in COVID-19 patients is important to develop countermeasures. Viral infections have previously been proposed to trigger autoimmune reactions, and it has been hypothesized that molecular mimicry plays a role in mediating autoimmune reactions.^7–9,11–13,29^ Here, a systematic analysis of gene-expression from cardiac tissues, based on bulk RNAseq data and single cell RNAseq data, combined with analysis of human genetic variation and SARS-CoV-2 genomes has led to the identification of candidate proteins and peptide regions therein that might be involved in immune cross reactivity (**Tables 2 and 3**). These newly identified identical peptides expand the set of known shared peptides between the human proteome and SARS-CoV-2, such as the furin cleavage site ‘RRARSVAS’ present in both human ENaC-α and the Spike glycoprotein.^10,30,31^ Further research is warranted to ascertain whether these mimicked peptides contribute to autoinflammatory pathology in the context of COVID-19 infection. Additionally, given the occurrence of myocarditis in some individuals shortly after receiving an mRNA COVID-19 vaccine, the potential for molecular mimicry between the antigen encoded by these vaccines (i.e., the pre-fusion stabilized Spike glycoprotein) and human cardiac proteins should be evaluated.^4–6,32,33^

There are a few limitations to this study. First, the human proteins that we surveyed were shortlisted based on their overexpression in cardiac tissue. There could be mimicked proteins that are shared between cardiac tissues and other tissues that are not accounted for in the current analysis. Second, there are other mechanisms that could contribute to autoimmunity after viral infection such as bystander activation, epitope spreading, and viral persistence^34^. Third, the presence of identical peptides in cardiac proteins and the SARS-CoV-2 proteome could occur due to chance. Comparing all SARS-CoV-2 8-mers to a set of brain enriched proteins and a set of skin enriched proteins shows a similar probability distribution of Hamming distance (**Supplementary Figure S2**), suggesting that the observed similarity with SARS-CoV-2 peptides is not specific to human cardiac proteins. Fourth, it is possible that peptides with lower degrees of similarity could contribute to immunologic mimicry, as T cells can be highly cross reactive against different major histocompatibility complex (MHC)-presented peptides.^35–39^

Taken together, by studying the intersection of human genetic variation in cardiac proteins and SARS-CoV-2 evolution, we have identified candidates of molecular mimicry that have potential to contribute to cardiac inflammation in the context of COVID-19. It will be important to perform follow-up functional studies evaluating the potential of SARS-CoV-2 reactive T cells and antibodies (e.g., from active or recovering COVID-19 patients) to cross-react with these peptides. Thus, we propose that SARS-CoV-2 variants harboring peptides identical to host heart-enriched proteins should be studied as ‘viral variants of cardiac interest’. We highlight that a similar strategy can be applied to identify and categorize plausible mimicry candidates from any human tissues that are targeted by other autoimmune responses in COVID-19 patients.

## Methods

### Identification of proteins enriched in cardiac tissue

Bulk RNA-sequencing (RNA-seq) data was accessed from the Genotype Tissue Expression (GTEx) project V8.^17^ For each sample, FASTQ files were processed using Salmon (in mapping-based mode) to quantify gene expression in transcripts per million (TPM). Specifically, the expression of each transcript isoform was first determined by passing FASTQ files to Salmon *quant* with the following parameters passed: validateMappings, rangeFactorizationBins 4, gcBias, biasSpeedSamp 10. All isoforms are then summed via a transcript-to-gene map, generating a gene-level expression value. GRCh38 was used as the reference, including cDNA and non-coding RNA.

For single cell RNA-seq studies, processed count matrices were accessed from Gene Expression Omnibus or other publicly available data repositories. There were two datasets analyzing heart tissues which captured cardiomyocytes, our main cell type of interest for this report.^20,40^ Other datasets captured a wide variety of immune, stromal, and parenchymal cell types from tissues including the respiratory tract, gastrointestinal tract, genitourinary tract, hepatobiliary system, skeletal muscle, brain, skin, eyes, and endocrine organs. Each dataset was processed using Scrublet and Seurat v3.0 as described previously.^18,41–43^ Cell type annotations were obtained from associated metadata files if available; otherwise, annotation was performed manually, guided by the cell types reported in the associated publication.

To identify genes that are overexpressed in cardiac tissue, we calculated fold change and Cohen’s D values between defined sample cohorts. For bulk RNA-seq data, Cohort A was defined as all GTEx heart samples (n = 861), and Cohort B was defined as all remaining GTEx samples except for those derived from skeletal muscle (n = 15,718). For single cell RNA-seq data, Cohort A was defined as all cells annotated as cardiomyocytes (n ∼ 8900 cells), and Cohort B was defined as all other cells from all processed studies (n ∼ 2.5 million cells). Fold change and Cohen’s D were calculated as follows:

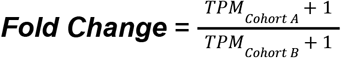

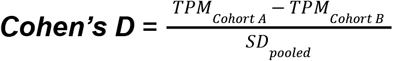, where the pooled standard deviation *SD*_*pooled*_ is defined as:

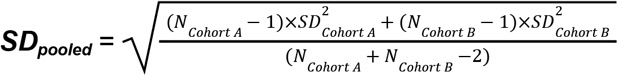, where N_cohort A_ and N_cohort B_ are the number of samples in Cohorts A and B, respectively, and SD_Cohort A_ and SD_Cohort B_ are the standard deviation of TPM values for the given gene in Cohorts A and B, respectively.

Genes with Fold Change ≥ 5 and Cohen’s D ≥ 0.5 from either the bulk or single cell RNA-seq analysis were considered to be enriched in cardiac tissue. In the volcano plots used to visualize these analyses, we filtered to genes with a TPM or CP10K value ≥ 1 in either Cohort A or Cohort B, and genes meeting the criteria for overexpression in heart or cardiomyocytes are colored in red.

For a control analysis, we also identified genes overexpressed in the brain or skin by bulk RNA-sequencing from the GTEx project. We used the same approach as described above, except that Cohort A was defined as either all brain samples (n = 2,351) or all skin samples (n = 1,305), and Cohort B was defined as all other samples.

### Comparison of 8-mers from reference sequences of cardiac proteins and SARS-CoV-2 proteins

The translated proteome from reference the SARS-CoV-2 genome (NC_045512.2) was downloaded from UniProt.^44,45^ A sliding window approach was used to enumerate all 8-mers from the 17 proteins in this viral proteome. Similarly, we used a sliding window approach to generate all 8-mers from the reference amino acid sequences of the previously defined 144 cardiac proteins, including the canonical isoforms and all described isoforms indicated in UniProt. We then performed a pairwise comparison of all 8-mers in these two groups by calculating the Hamming distance using the *stringdist* function from the stringdist package (version 0.9.8) in R (version 4.0.3). In a control analysis, we used the same approach to calculate the Hamming distance between all SARS-CoV-2 8-mers and the control sets of 369 human proteins enriched in the brain or 198 human proteins enriched in the skin (described above).

### Assessing the impact of human and SARS-CoV-2 variants on cardiac peptide matches

To assess the impact of human genetic variation on potential molecular mimicry, we retrieved all missense variants from the gnomAD database for the previously identified cardiac proteins that had at least one 8-mer similar to peptide in the SARS-CoV-2 reference proteome (Hamming distance = 1).^22^ We used the gnomad-api (https://gnomad.broadinstitute.org/api) to fetch the variant calls from gnomad_r2_1 version from the Human GRCh37 genome assembly. The variants in this gnomad version (GRCh37/hg19) are derived from 125,748 exome sequences and 15,708 whole-genome sequences from unrelated individuals sequenced as part of various disease-specific and population genetic studies. For any variants that alter the amino acid sequence of a potentially mimicked peptide, we determined whether the mutation resulted in an exact match (Hamming distance = 0) to the corresponding 8-mer from the SARS-CoV-2 reference proteome.

To assess the impact of viral evolution on potential molecular mimicry, we queried the cardiac 8-mers with Hamming distance of 1 (including any alterations of these 8-mers arising from human genetic variation as described above) against all protein variants encoded in 4,854,709 SARS-CoV-2 genomes deposited in the GISAID database (last accessed 11/5/2021).^24^ Here, we determined whether any mutations in viral genomes (relative to the reference sequence) resulted in 8-mers which exactly match one or more cardiac peptides.

### Evaluation of mimicked peptides for inclusion in immune epitopes

For any 8-mers which showed an exact match between a cardiac peptide (reference or variant sequences) and a SARS-CoV-2 peptide (reference or variant sequences), we queried the 8-mer using the Immune Epitope Database (IEDB; www.iedb.org) and Analysis Resource.^23^

We searched for any linear peptide epitope with a Blast similarity of at least 90% from any human host that had positive experimental evidence in any assay (T cell, B cell, or MHC Ligand). No MHC class restrictions or disease filters were applied.

## Supplementary Material

**Supplementary Figure S1:**
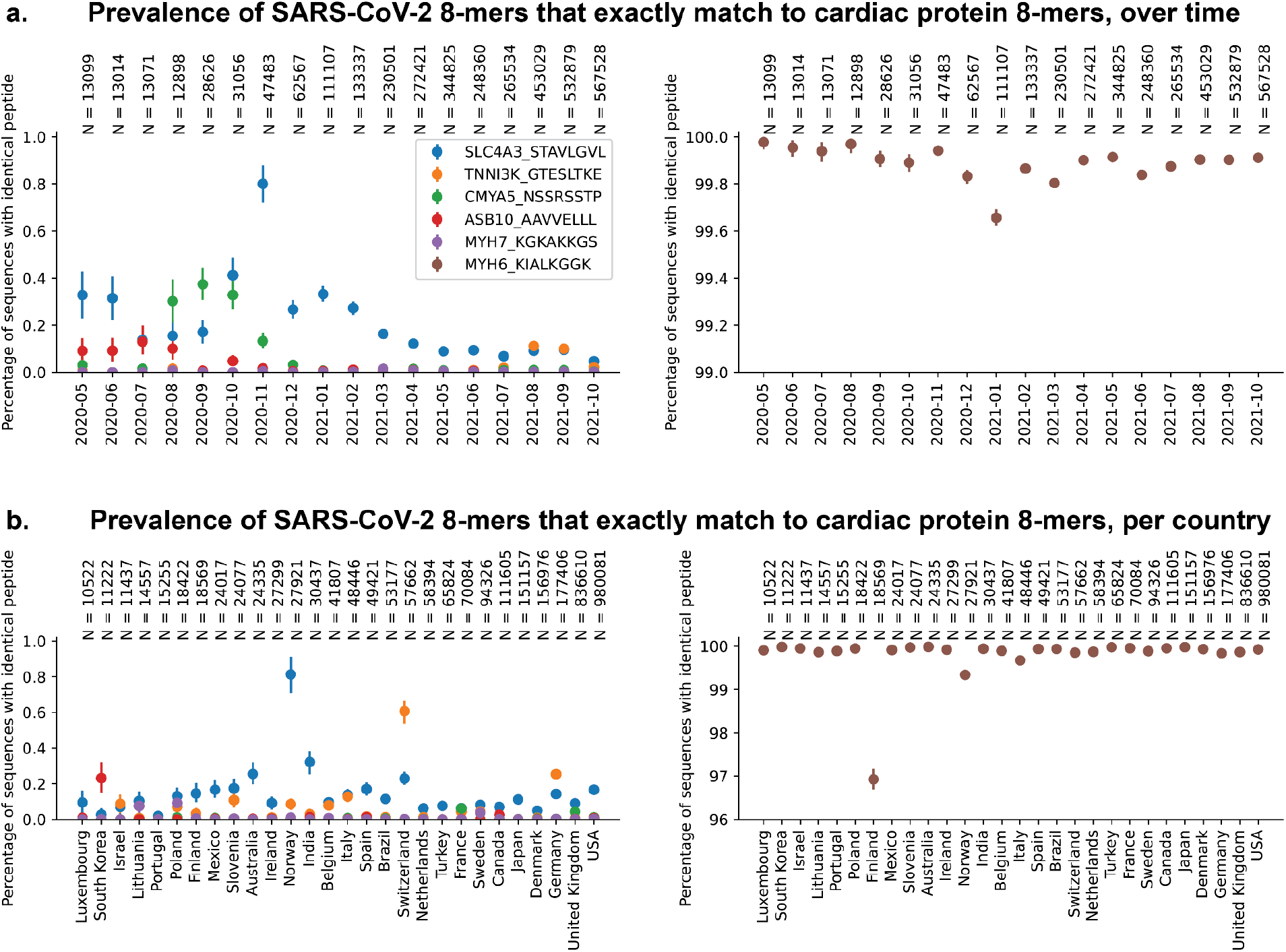
Prevalence of SARS-CoV-2 variants with exact 8-mer matches to human cardiac proteins. Prevalence is reported as the percentage of SARS-CoV-2 sequences with exact 8-mer matches during 1-month time intervals (**a**) or across all sequences reported by a country (**b**). Prevalence of the wild-type SARS-CoV-2 8-mer that exactly matches with the MYH6 peptide, described in the main text, is shown separately on the right (brown). Data is only shown for months and countries in which more than 10,000 SARS-CoV-2 genomes were reported to GISAID.

**Supplementary Figure S2.**
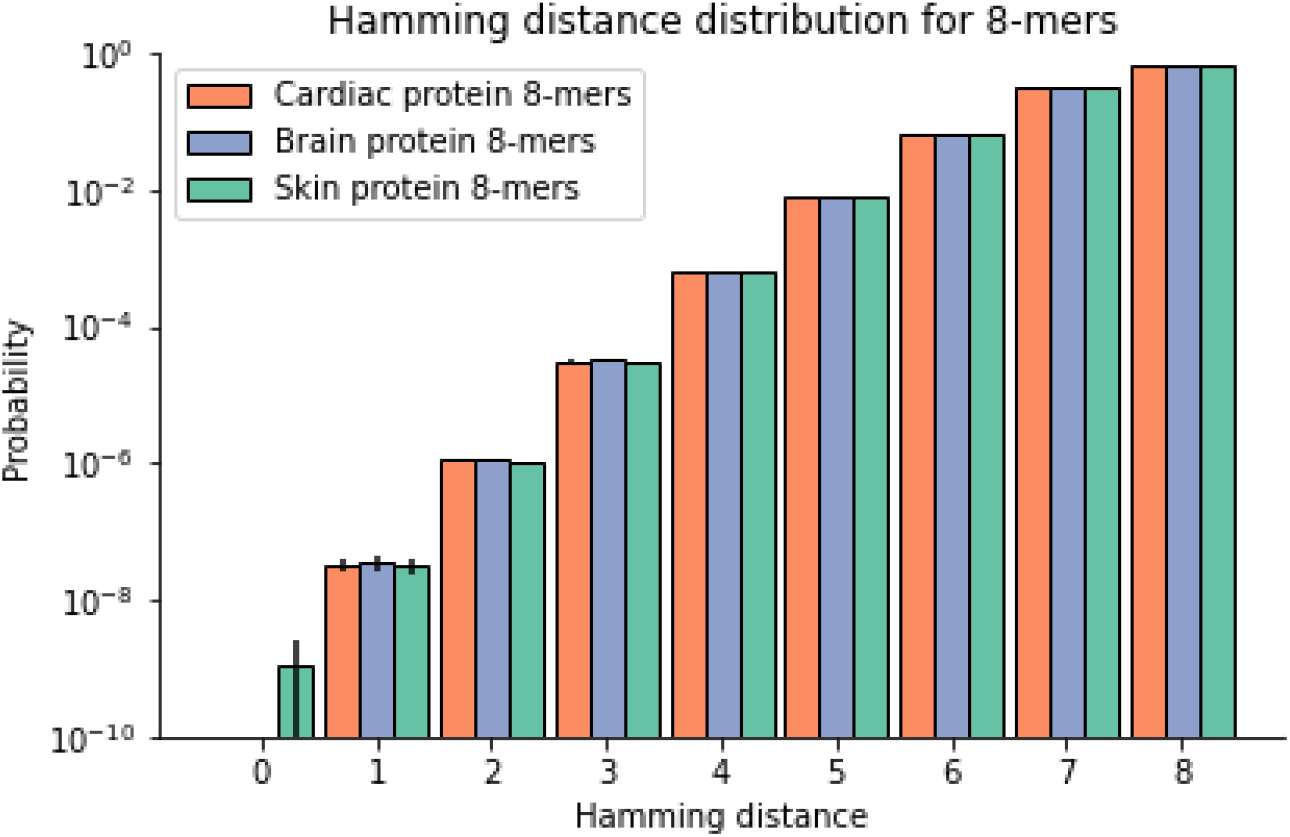
Observed probability distribution for the Hamming distance between all 8-mers in the SARS-CoV-2 proteome and all 8-mers in reference sets of human proteins. Shown is the distribution of Hamming distance between all SARS-CoV-2 linear 8-mer peptides (n=9,926 peptides) and all linear 8-mer peptides from canonical isoforms of human cardiac proteins (orange, n=129,415 peptides), brain proteins (blue, n=241,046 peptides), and skin proteins (green, n=88,628 peptides). Error bars represent 95% confidence intervals.

**Supplementary Table S1.**
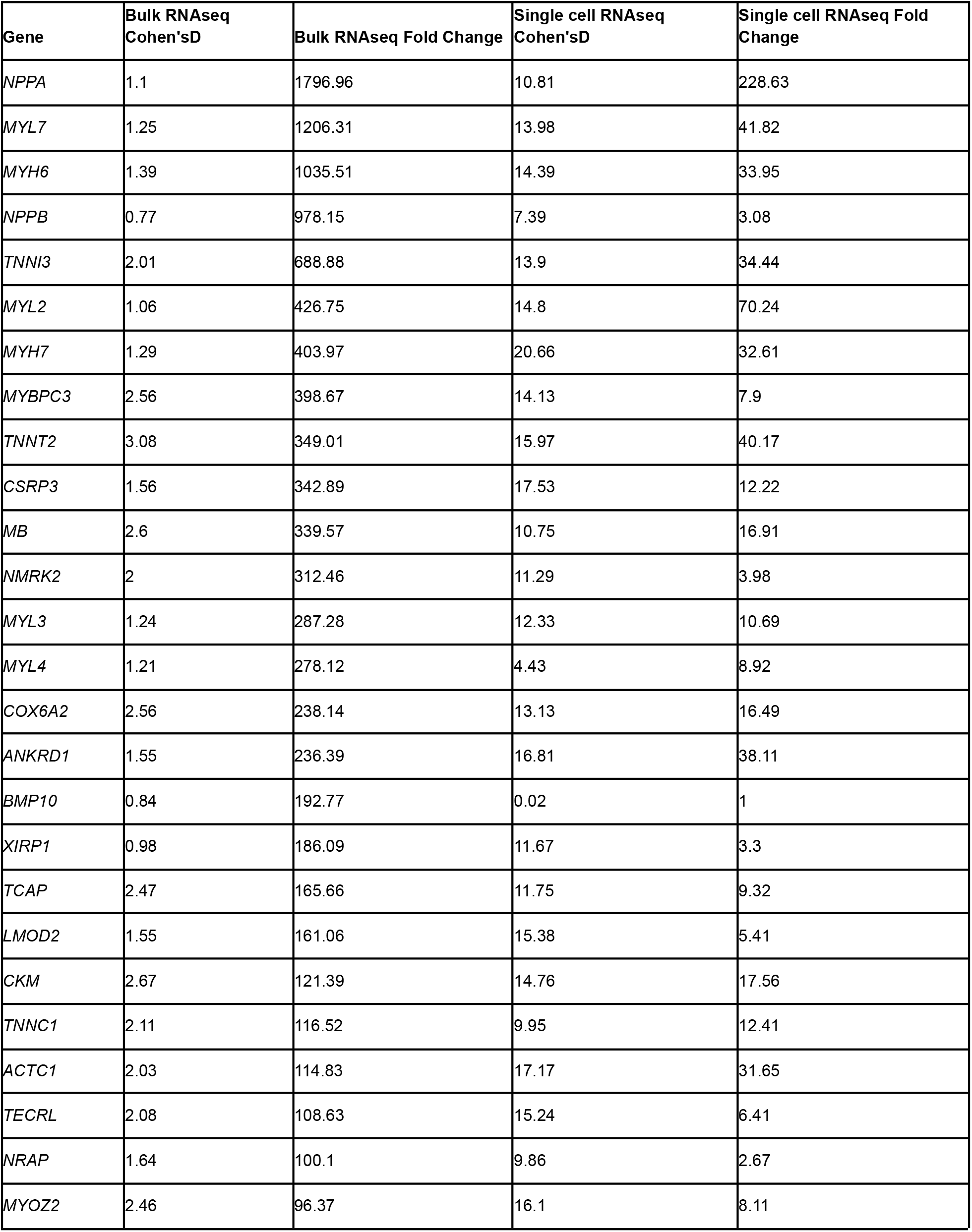
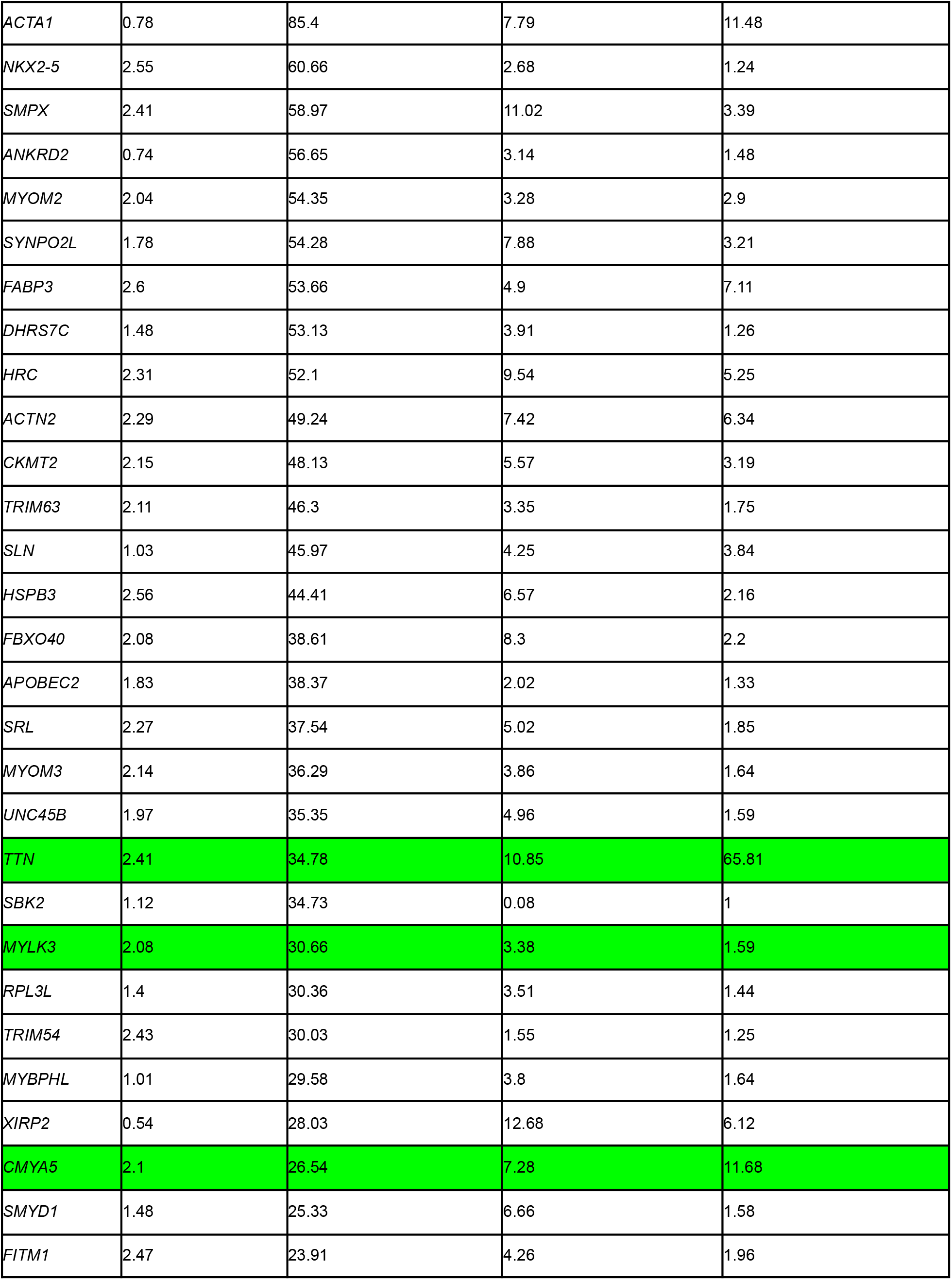
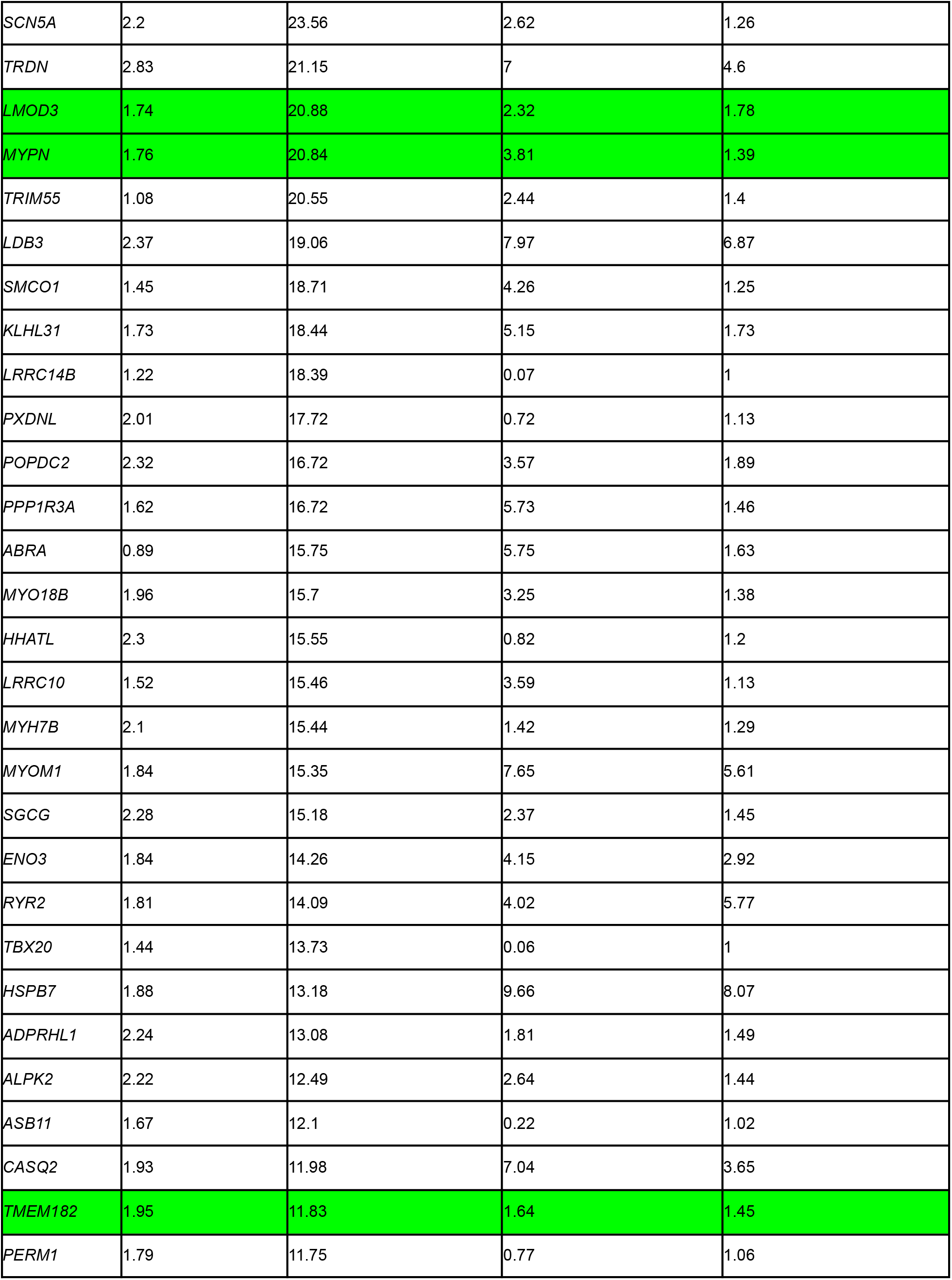
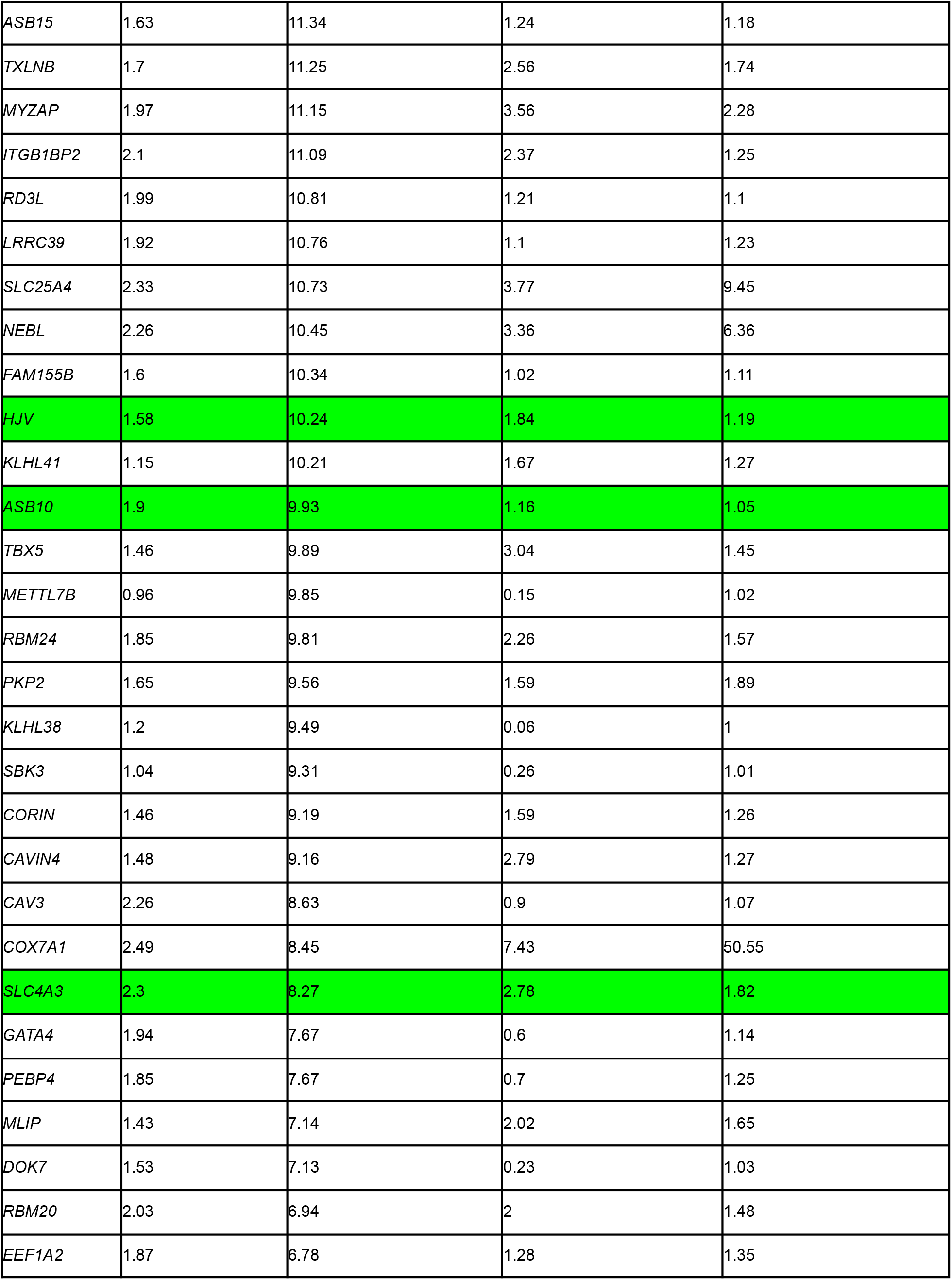
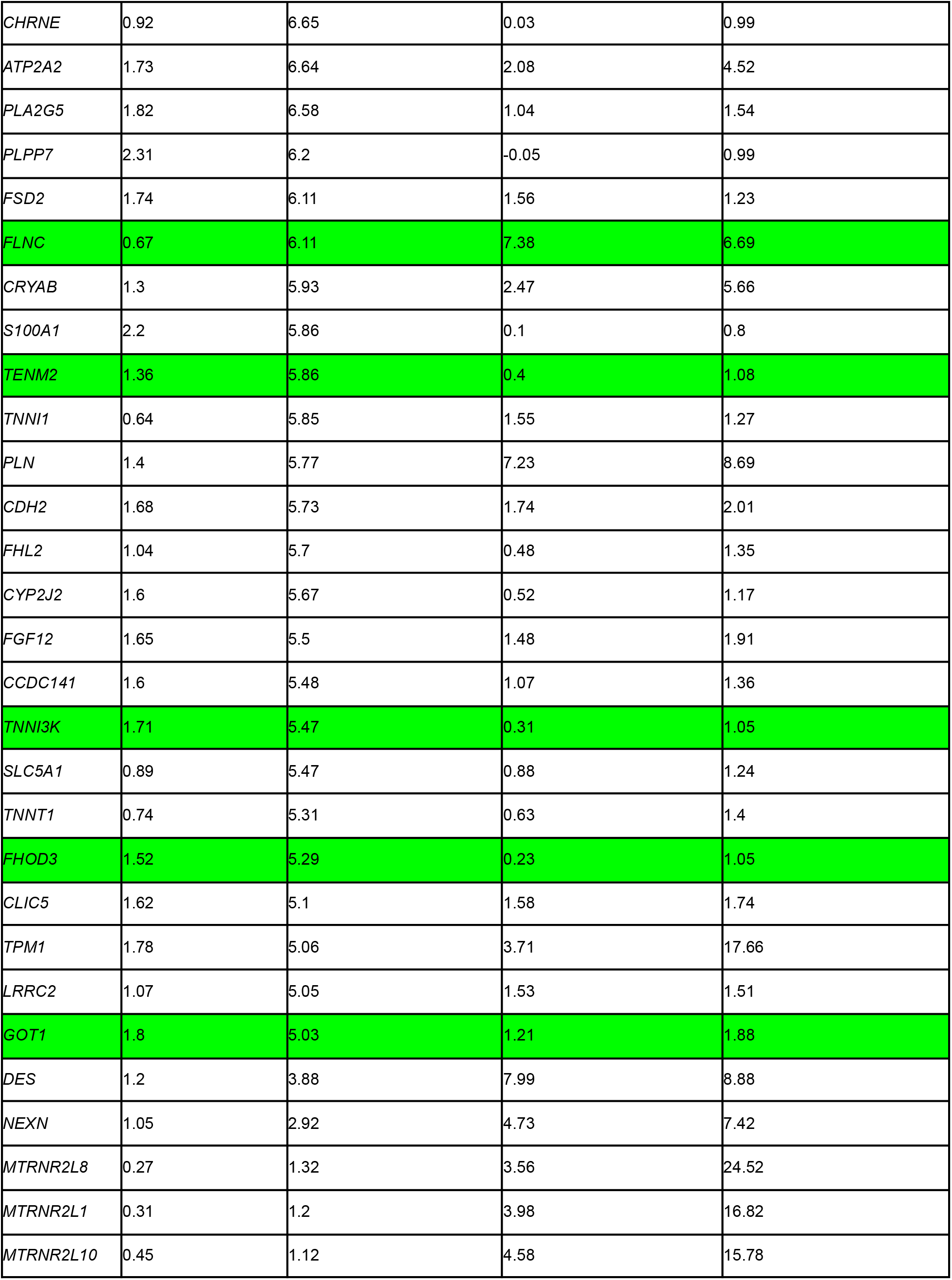
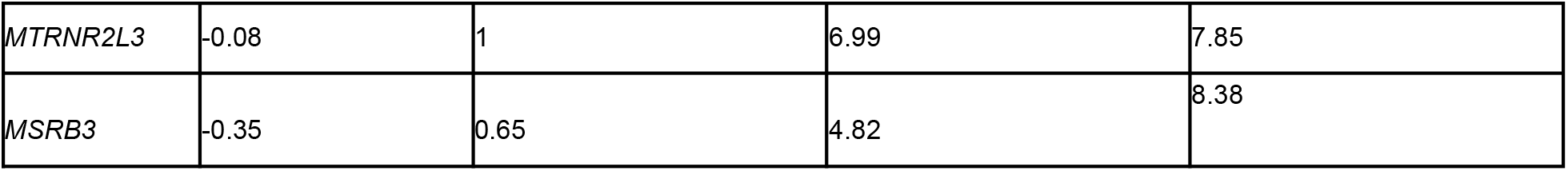
List of human cardiac proteins identified based on Bulk RNA-seq and Single cell RNA-seq. Rows with human genes from Tables 2 and 3 are highlighted.

## References

1. Chung, M. K. et al. COVID-19 and Cardiovascular Disease: From Bench to Bedside. Circ. Res. 128, 1214–1236 (2021).

2. Puntmann, V. O. et al. Outcomes of Cardiovascular Magnetic Resonance Imaging in Patients Recently Recovered From Coronavirus Disease 2019 (COVID-19). JAMA Cardiol 5, 1265–1273 (2020).

3. Barda, N. et al. Safety of the BNT162b2 mRNA Covid-19 Vaccine in a Nationwide Setting. N. Engl. J. Med. 385, 1078–1090 (2021).

4. CDC. Myocarditis and Pericarditis After mRNA COVID-19 Vaccination. https://www.cdc.gov/coronavirus/2019-ncov/vaccines/safety/myocarditis.html (2021).

5. Witberg, G. et al. Myocarditis after Covid-19 Vaccination in a Large Health Care Organization. New England Journal of Medicine (2021) doi:10.1056/nejmoa2110737.

6. Simone, A. et al. Acute Myocarditis Following COVID-19 mRNA Vaccination in Adults Aged 18 Years or Older. JAMA Intern. Med. (2021) doi:10.1001/jamainternmed.2021.5511.

7. Bozkurt, B., Kamat, I. & Hotez, P. J. Myocarditis With COVID-19 mRNA Vaccines. Circulation 144, 471–484 (2021).

8. Proal, A. D. & VanElzakker, M. B. Long COVID or Post-acute Sequelae of COVID-19 (PASC): An Overview of Biological Factors That May Contribute to Persistent Symptoms. Front. Microbiol. 12, 698169 (2021).

9. Galeotti, C. & Bayry, J. Autoimmune and inflammatory diseases following COVID-19. Nature reviews. Rheumatology vol. 16 413–414 (2020).

10. Kanduc, D. From Anti-SARS-CoV-2 Immune Responses to COVID-19 via Molecular Mimicry. Antibodies (Basel) 9, (2020).

11. Gowthaman, U. & Eswarakumar, V. P. Molecular mimicry: good artists copy, great artists steal. Virulence 4, 433–434 (2013).

12. Cusick, M. F., Libbey, J. E. & Fujinami, R. S. Molecular mimicry as a mechanism of autoimmune disease. Clin. Rev. Allergy Immunol. 42, 102–111 (2012).

13. Oldstone, M. B. A. Molecular Mimicry, Microbial Infection, and Autoimmune Disease: Evolution of the Concept. in Molecular Mimicry: Infection-Inducing Autoimmune Disease (ed. Oldstone, M. B. A.) 1–17 (Springer Berlin Heidelberg, 2005).

14. Adderson, E. E., Shikhman, A. R., Ward, K. E. & Cunningham, M. W. Molecular analysis of polyreactive monoclonal antibodies from rheumatic carditis: human anti-N-acetylglucosamine/anti-myosin antibody V region genes. J. Immunol. 161, 2020–2031 (1998).

15. Cunningham, M. W., Antone, S. M., Smart, M., Liu, R. & Kosanke, S. Molecular analysis of human cardiac myosin-cross-reactive B- and T-cell epitopes of the group A streptococcal M5 protein. Infect. Immun. 65, 3913–3923 (1997).

16. Cunningham, M. W. Autoimmunity and molecular mimicry in the pathogenesis of post-streptococcal heart disease. Front. Biosci. 8, s533–43 (2003).

17. GTEx Consortium. The Genotype-Tissue Expression (GTEx) project. Nat. Genet. 45, 580–585 (2013).

18. Venkatakrishnan, A. J. et al. Knowledge synthesis of 100 million biomedical documents augments the deep expression profiling of coronavirus receptors. Elife 9, (2020).

19. Rozenblatt-Rosen, O., Stubbington, M. J. T., Regev, A. & Teichmann, S. A. The Human Cell Atlas: from vision to reality. Nature 550, 451–453 (2017).

20. Han, X. et al. Construction of a human cell landscape at single-cell level. Nature 581, 303–309 (2020).

21. UniProt Consortium. UniProt: the universal protein knowledgebase in 2021. Nucleic Acids Res. 49, D480–D489 (2021).

22. Karczewski, K. J. et al. The mutational constraint spectrum quantified from variation in 141,456 humans. Nature 581, 434–443 (2020).

23. Vita, R. et al. The Immune Epitope Database (IEDB): 2018 update. Nucleic Acids Res. 47, D339–D343 (2019).

24. Shu, Y. & McCauley, J. GISAID: Global initiative on sharing all influenza data - from vision to reality. Euro Surveill. 22, (2017).

25. IEDB.org: Free epitope database and prediction resource. https://www.iedb.org/home_v3.php.

26. Thorsen, K. et al. Loss-of-activity-mutation in the cardiac chloride-bicarbonate exchanger AE3 causes short QT syndrome. Nat. Commun. 8, 1696 (2017).

27. Anion exchange protein 3. https://www.uniprot.org/uniprot/P48751.

28. Altschul, S. F., Gish, W., Miller, W., Myers, E. W. & Lipman, D. J. Basic local alignment search tool. Journal of Molecular Biology vol. 215 403–410 (1990).

29. Ehrenfeld, M. et al. Covid-19 and autoimmunity. Autoimmunity Reviews vol. 19 102597 (2020).

30. Anand, P., Puranik, A., Aravamudan, M., Venkatakrishnan, A. J. & Soundararajan, V. SARS-CoV-2 strategically mimics proteolytic activation of human ENaC. Elife 9, (2020).

31. Venkatakrishnan, A. J. et al. Benchmarking evolutionary tinkering underlying human-viral molecular mimicry shows multiple host pulmonary-arterial peptides mimicked by SARS-CoV-2. Cell Death Discov 6, 96 (2020).

32. Corbett, K. S. et al. SARS-CoV-2 mRNA vaccine design enabled by prototype pathogen preparedness. Nature 586, 567–571 (2020).

33. Walsh, E. E. et al. Safety and Immunogenicity of Two RNA-Based Covid-19 Vaccine Candidates. N. Engl. J. Med. 383, 2439–2450 (2020).

34. Fujinami, R. S., von Herrath, M. G., Christen, U. & Whitton, J. L. Molecular mimicry, bystander activation, or viral persistence: infections and autoimmune disease. Clin. Microbiol. Rev. 19, 80–94 (2006).

35. Sewell, A. K. Why must T cells be cross-reactive? Nat. Rev. Immunol. 12, 669–677 (2012).

36. Borbulevych, O. Y., Santhanagopolan, S. M., Hossain, M. & Baker, B. M. TCRs used in cancer gene therapy cross-react with MART-1/Melan-A tumor antigens via distinct mechanisms. J. Immunol. 187, 2453–2463 (2011).

37. Scott, D. R., Borbulevych, O. Y., Piepenbrink, K. H., Corcelli, S. A. & Baker, B. M. Disparate degrees of hypervariable loop flexibility control T-cell receptor cross-reactivity, specificity, and binding mechanism. J. Mol. Biol. 414, 385–400 (2011).

38. Garcia, K. C. et al. Structural basis of plasticity in T cell receptor recognition of a self peptide-MHC antigen. Science 279, 1166–1172 (1998).

39. Wooldridge, L. et al. A single autoimmune T cell receptor recognizes more than a million different peptides. J. Biol. Chem. 287, 1168–1177 (2012).

40. Wang, L. et al. Single-cell reconstruction of the adult human heart during heart failure and recovery reveals the cellular landscape underlying cardiac function. Nat. Cell Biol. 22, 108–119 (2020).

41. Doddahonnaiah, D. et al. A Literature-Derived Knowledge Graph Augments the Interpretation of Single Cell RNA-seq Datasets. Genes 12, (2021).

42. Stuart, T. et al. Comprehensive Integration of Single-Cell Data. Cell 177, 1888–1902.e21 (2019).

43. Wolock, S. L., Lopez, R. & Klein, A. M. Scrublet: Computational identification of cell Doublets in Single-cell transcriptomic data. Cell Syst. 8, 281–291.e9 (2019).

44. UniProt. https://covid-19.uniprot.org/uniprotkb?query=+&facets=other_organism:Severe%20acute%20respiratory%20syndrome%20coronavirus%202.

45. Severe acute respiratory syndrome coronavirus 2 isolate Wuhan-Hu-1, co - Nucleotide - NCBI. https://www.ncbi.nlm.nih.gov/nuccore/NC_045512.2.

